# Cancer-associated fibroblasts exert a pro-angiogenic activity in Merkel cell carcinoma

**DOI:** 10.1101/2021.07.30.454425

**Authors:** Silvia Albertini, Licia Martuscelli, Cinzia Borgogna, Sanamjeet Virdi, Daniela Indenbirken, Irene Lo Cigno, Gloria Griffante, Federica Calati, Renzo Boldorini, Nicole Fischer, Marisa Gariglio

**Affiliations:** Institute for Medical Microbiology, Virology and Hygiene, University Medical Center Hamburg-Eppendorf, Hamburg, 20251, Germany; Virology Unit, Department of Translational Medicine, Novara Medical School, UPO, Novara, 28100, Italy; Leibniz Institute for Experimental Virology, Technology Platform Next Generation Sequencing, Hamburg, 20251, Germany; Pathology Unit, Department of Health Sciences, Novara Medical School, UPO, Novara, 28100, Italy

**Keywords:** angiogenesis, cancer associated fibroblasts, Merkel cell carcinoma, Merkel cell polyomavirus, tumor microenvironment

## Abstract

The tumor microenvironment (TME) is a complex niche enveloping a tumor formed by extracellular matrix, blood vessels, immune cells, and fibroblasts constantly interacting with cancer cells. Although TME is increasingly recognized as a major player in cancer initiation and progression in many tumor types, its involvement in Merkel cell carcinoma (MCC) pathogenesis is currently unknown. Here, we provide the first molecular and functional characterization of cancer-associated fibroblasts (CAFs), the major TME component, in MCC patient-derived xenografts. We show that subcutaneous co-injection of patient-derived CAFs and human MCC MKL-1 cells into SCID mice significantly promotes tumor growth and metastasis. These fast-growing xenografts are characterized by areas densely populated with human CAFs, mainly localized around blood vessels. We also provide evidence that the growth-promoting activity of MCC-derived CAFs is mediated by the APA/Ang II-III/AT_1_R axis, with the expression of aminopeptidase A (APA) in CAFs being the upstream triggering event. Altogether, our findings point to APA as a potential marker for MCC prognostic stratification and a novel candidate therapeutic target.

## Introduction

Merkel cell carcinoma (MCC) is a highly aggressive skin cancer with an increased mortality rate compared to stage-matched melanoma. MCC mainly affects older and immunosuppressed patients who, at the time of diagnosis, often present with advanced clinical stage disease (Becker *et al*, 2017). The 5-year survival rate is dependent on the stage of metastasis and is approximately 51% in patients with a localized tumor, 35% in cases with loco regional metastasis, and as low as 15% in those with distant metastasis (Harms *et al*, 2016; Trinidad *et al*, 2019). The majority of MCCs, approximately 80% in the Northern Hemisphere, is caused by clonal integration of Merkel cell polyomavirus (MCPyV) into the host genome with persistent expression of the viral oncoproteins, small T-Antigen (sT) and large T-Antigen (LT) (Feng *et al*, 2008).

While virus positive (+) MCCs have relatively few genomic aberrations in the host genome and exhibit few mutations, virus negative (-) MCCs, on the other hand, have a high number of genomic aberrations and a predominant ultraviolet (UV) mutational signature (Goh *et al*, 2016; Gonzalez-Vela *et al*, 2017; Harms *et al*., 2016; Harms *et al*, 2015). Despite these notable differences in MCC etiology, both virus + and virus - MCCs are similar in presentation, prognosis, and response to therapy (Gonzalez-Vela *et al*., 2017).

MCPyV belongs to the family *Polyomaviridae*, which are highly prevalent, non-enveloped double-stranded DNA viruses causing opportunistic infections. MCPyV is the only human family member causally linked to tumorigenesis. Virus + MCCs show monoclonally-integrated replication-defective forms of the viral genome with tumor hallmark mutations in the viral early gene region (Feng *et al*., 2008). Although MCPyV is clearly the etiologic agent of virus + MCCs, the correlations between the molecular phenotype, therapeutic approaches, and clinical outcome of MCPyV+ MCC remain poorly understood.

It is well established that tumor growth and progression are profoundly influenced by the tumor microenvironment (TME), which includes fibroblasts, inflammatory cells, blood vessels, extracellular matrix (ECM), and basement membranes (BMs) (Hinshaw & Shevde, 2019). Specifically, the tumor populating fibroblasts—termed cancer-associated fibroblasts (CAFs)—are desmoplastic cells that, once activated, contribute to cancer development through a variety of mechanisms. These include the release of various tumor-promoting factors, such as cytokines and chemokines, which, in turn, foster cancer cell growth and angiogenesis. CAFs are spindle-shaped, non-epithelial and non-immune cells embedded in the ECM that can be easily propagated in adherent cell cultures (Kalluri, 2016). Many reports have already extensively demonstrated that the desmoplastic stroma—mostly sustained by CAFs—in colorectal, breast, ovarian, head and neck cancers is associated with poor prognosis (Heichler *et al*, 2020; Karakasheva *et al*, 2018; Leung *et al*, 2018; Wen *et al*, 2019).

Despite the central role of CAFs in carcinogenesis, the function of CAFs in MCC pathogenesis and disease progression has been poorly investigated. Few publications on tumor infiltrating immune cells suggest a correlation between lymphocyte density, activation state, genetic diversity, and improved survival of MCC patients (Mendoza *et al*, 2020; Paulson *et al*, 2014; Samimi *et al*, 2019). A more recent study provided evidence of exosome transmitted cargo on the TME in MCC. Exosome transmitted miRNA-375 induced activation of αSMA-1, CXCL2, and IL- 1β genes, suggesting that horizontally transferred miR-375 is critical for polarizing fibroblasts toward a CAF phenotype in MCC (Fan *et al*, 2021).

To gain further insight into the functional role of the TME in MCC carcinogenesis, we have characterized the mechanism and functional role of MCC-derived CAFs – the major component of TME – in tumor growth and angiogenesis. Applying immunohistochemistry, transcriptomics, and functional assays, including tube formation and contraction assays, and in vivo tumorigenic assays, we show that patient-derived CAFs can differentially support MCC growth and angiogenesis through the renin-angiotensin system (RAS), specifically the APA/Ang II-III/AT_1_R axis, ultimately leading to increased angiogenesis and tumor progression. We also show that aminopeptidase A (APA) expression mainly occurs in fibroblasts, indicating a prominent role of CAFs in fostering a pro-angiogenic tumor microenvironment.

Overall, our findings provide the rationale for further evaluation of APA expression and function in MCC carcinogenesis as a potential marker of CAF-activated phenotype, which in turn may allow prognostic stratification of disease progression. In addition, pharmaceutical manipulation of this pathway may help develop future therapies against MCC.

## Results

### Characterization of MCC patients and corresponding CAFs

We collected fresh tumor tissues from 9 MCC patients (Pts1-9) who underwent surgical excision of 3 primary tumors (pPts 4, 8 and 9), 5 local recurrences (rPts 1, 2, 3, 5, 7), and 1 lymph node metastasis (nPt 6). The clinicopathological features of the study cohort are summarized in Table 1. Screening for MCPyV by both immunohistochemistry (IHC) (LT-antigen expression) and PCR (viral DNA detection) revealed that 6 MCCs were MCPyV+ (Pts 1, 2, 5, 6, 8, and 9), with viral loads ranging from 2 to 12 copies/cell (Table 1). In contrast, MCCs from Pts 3, 4, and 7 were MCPyV-. In a median follow-up period of 30 months, one patient, Pt3, died from MCC brain metastasis, three months after surgical excision of a relapsed MCC.

**Table 1.**
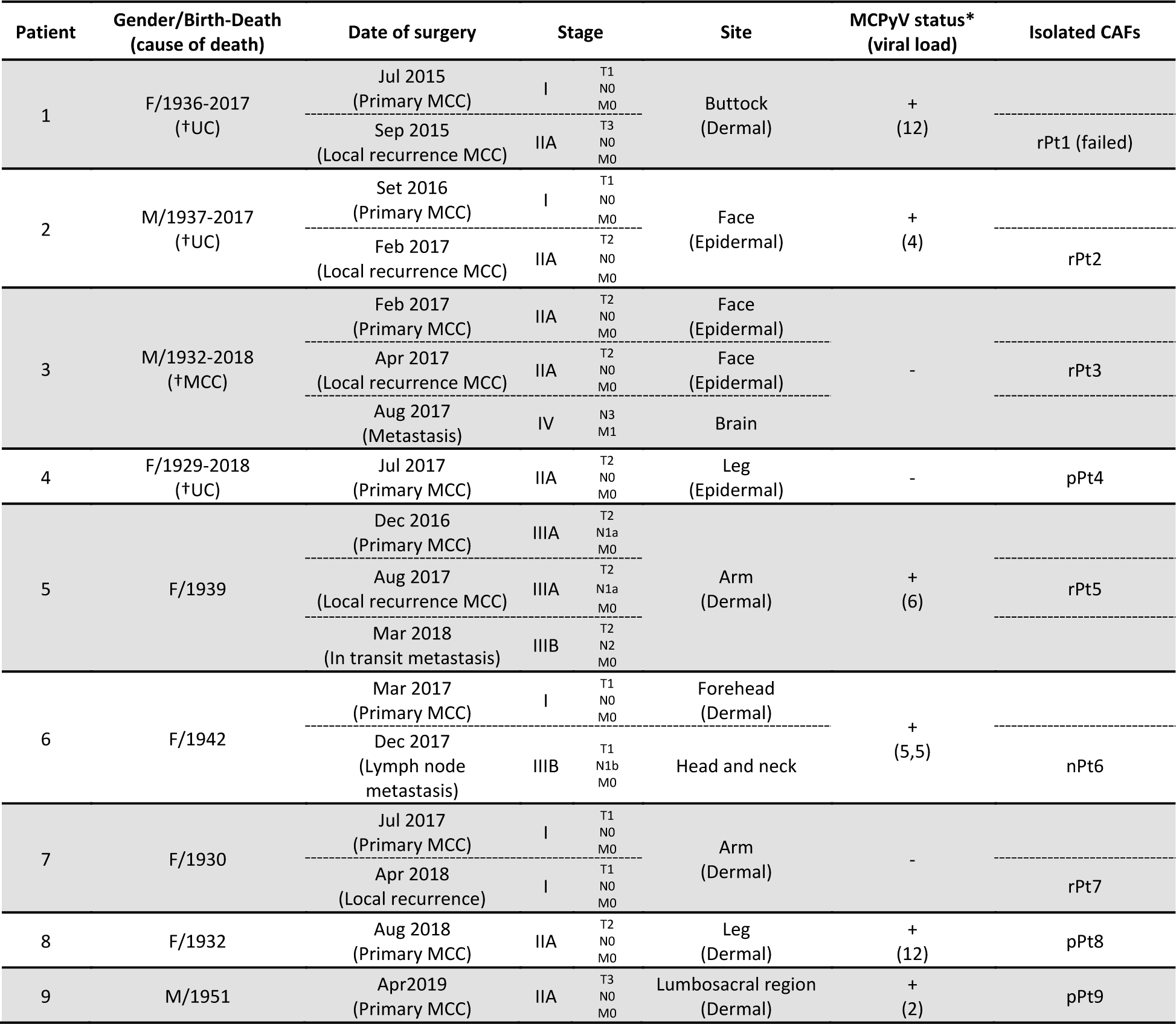
Clinicopathological features of the MCC cohort.

CAFs were successfully isolated from 8 of the aforementioned 9 MCC patients (Pts 2-9) by mechanic dissociation of the tumor masses. CAFs were named according to the stage of the relative tumor: primary (pCAFs); relapsed (rCAFs); or lymph node metastasis (nCAFs).

All patient-derived CAFs expressed, albeit to different extents, αSMA-1 and vimentin, hallmarks of activated fibroblasts and mesenchymal origin. In addition, CAFs proved negative for CD31 or cytokeratin 14 expression, excluding contamination by endothelial or epithelial cells (Figure 1A). In addition, all patient-derived CAFs, except for nPt6, showed enhanced extracellular matrix contractility compared to that of HFFs, as judged by an *in vitro* collagen contraction assay (Figure 1B). We next evaluated the mRNA expression levels of pro-inflammatory cytokines and chemokines known to be frequently upregulated in CAFs isolated from other carcinoma types (Kalluri, 2016). RT-qPCR revealed that IL-8, IL-6, CXCL12, and TGF-β1 were generally upregulated in MCC-derived CAFs when compared to normal human foreskin fibroblasts (HFFs) (Figure 1C), with few notable exceptions: IL-8 in pPt8 and pPt9, and IL-6, CXCL12, and TGF-β1 in pPt9.

**Figure 1.**
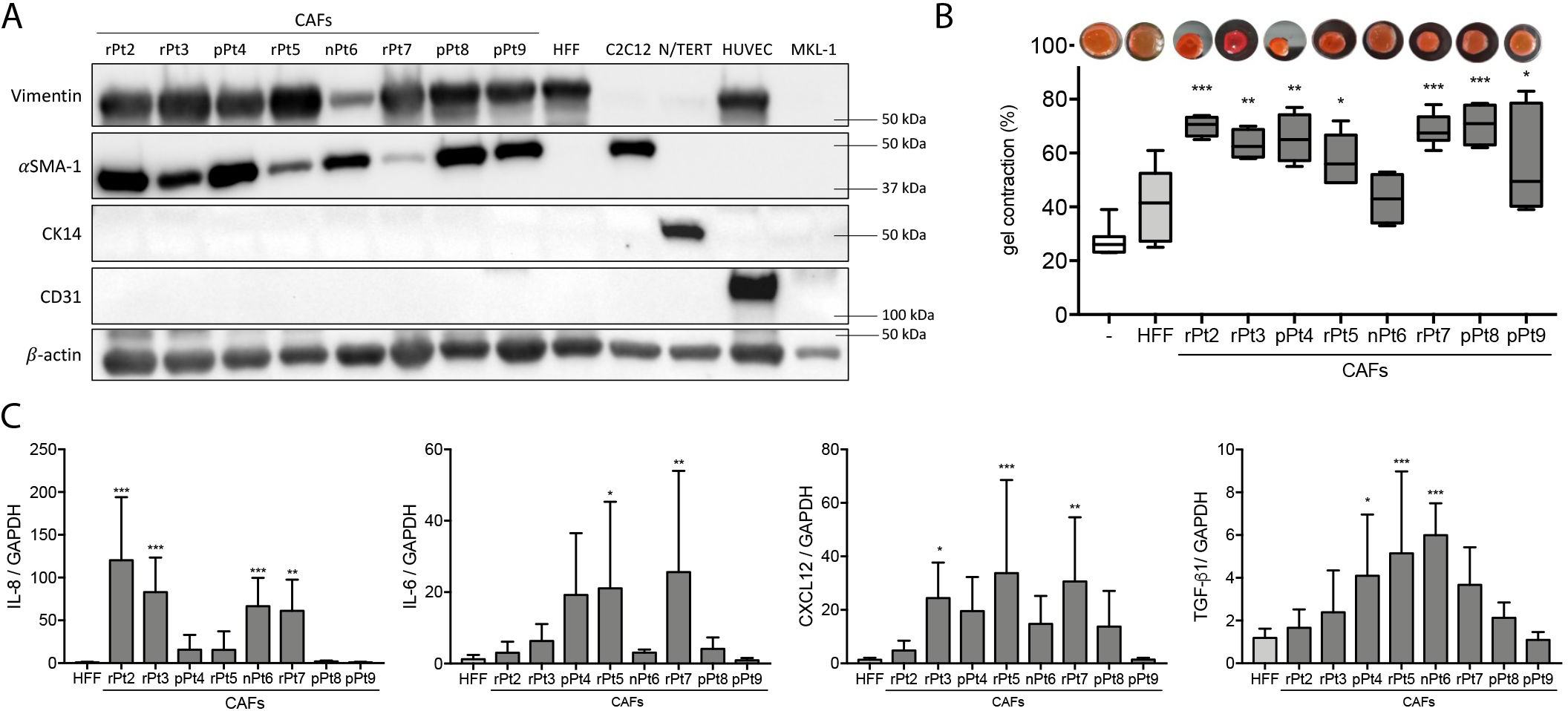
Characterization of MCC-derived cancer associated fibroblasts (CAFs). **A** Immunoblot analysis on total cell lysates from the isolated CAFs along with a panel of control cell lines [human foreskin fibroblasts (HFFs); mouse myofibroblasts, C2C12; immortalized human keratinocytes, N/TERT; human umbilical vein endothelial cells (HUVECs)] probed with antibodies against vimentin, αSMA-1, CK14, CD31, and β-actin. Representative blots from triplicate experiments are shown. **B** Collagen contractility assay evaluated as percentage of gel contraction in gel without cells (-), and gel containing normal fibroblasts (HFFs) or individual CAFs (rPt2-pPt9); representative pictures are shown on the top of the box plot graph (*n* = 3). **C** qRT-PCR analysis for IL-8, IL-6, CXCL12, and TGF-β1 mRNA expression levels in total RNA extracts from CAFs or HFFs. Values were normalized to GAPDH mRNA and plotted as fold induction over HFFs (*n* = 4). Data Information: In (B), the whisker-box plots represent the 25^th^-75^th^ percentiles, with midlines indicating the median values and whiskers extending to the minimum and maximum values. In (C), values are represented as mean ± SD. *n* indicates the number of independent experiments. In (B-C), the significance levels are as follows: \**P* < 0.05, \*\**P* < 0.01, \*\*\**P* < 0.001 *vs* HFFs; classical one- way ANOVA followed by post-hoc Bonferroni correction.

To further characterize the stromal components of the original tumors, formalin fixed paraffin embedded (FFPE) tissue sections derived from primary and relapsed MCCs—if available—were analyzed by IHC using anti-vimentin (anti-Vim) staining. The overall density of Vim^+^ cells varied significantly between samples, ranging from 10% to 60% [see Figure 2A for representative panels of low density (pPt5) and high density (pPt6) stromal component]. Interestingly, we observed a slight increase in Vim^+^ stromal components in the relapsed MCCs compared to the primary tumors (Figure 2B, grey pie chart), which led us to assess the localization of αSMA-1^+^ cells in the tumor stroma. Given the established regulatory association between CAFs and immune cells (Kalluri, 2016), we further aimed to determine the localization of CD8^+^ cytotoxic T cells and CD68^+^ macrophages in the desmoplastic stroma. As shown in Figure 2 and Figure EV1, αSMA-1^+^ cells were mainly localized in the stromal strakes and tumor edges, with only a few αSMA-1^+^, CD8^+^, and CD68^+^ cells penetrating the tumor nests. Within the stromal strakes, only the primary and relapsed specimens from Pt5 (Figure 2B) showed a higher proportion of CD8^+^ (red) and CD68^+^ (blue) cells over a background of more scattered αSMA-1^+^ cells (green). A substantial number of CD68^+^ was also observed in Pt2, pPt3, and pPt9. In all the other sections, the overall number of αSMA-1^+^ cells was higher and, for the most part, overlapped with the vimentin staining. Overall, the MCC stroma shows considerable variability in terms of cellular composition, density, and/or localization.

**Figure 2.**
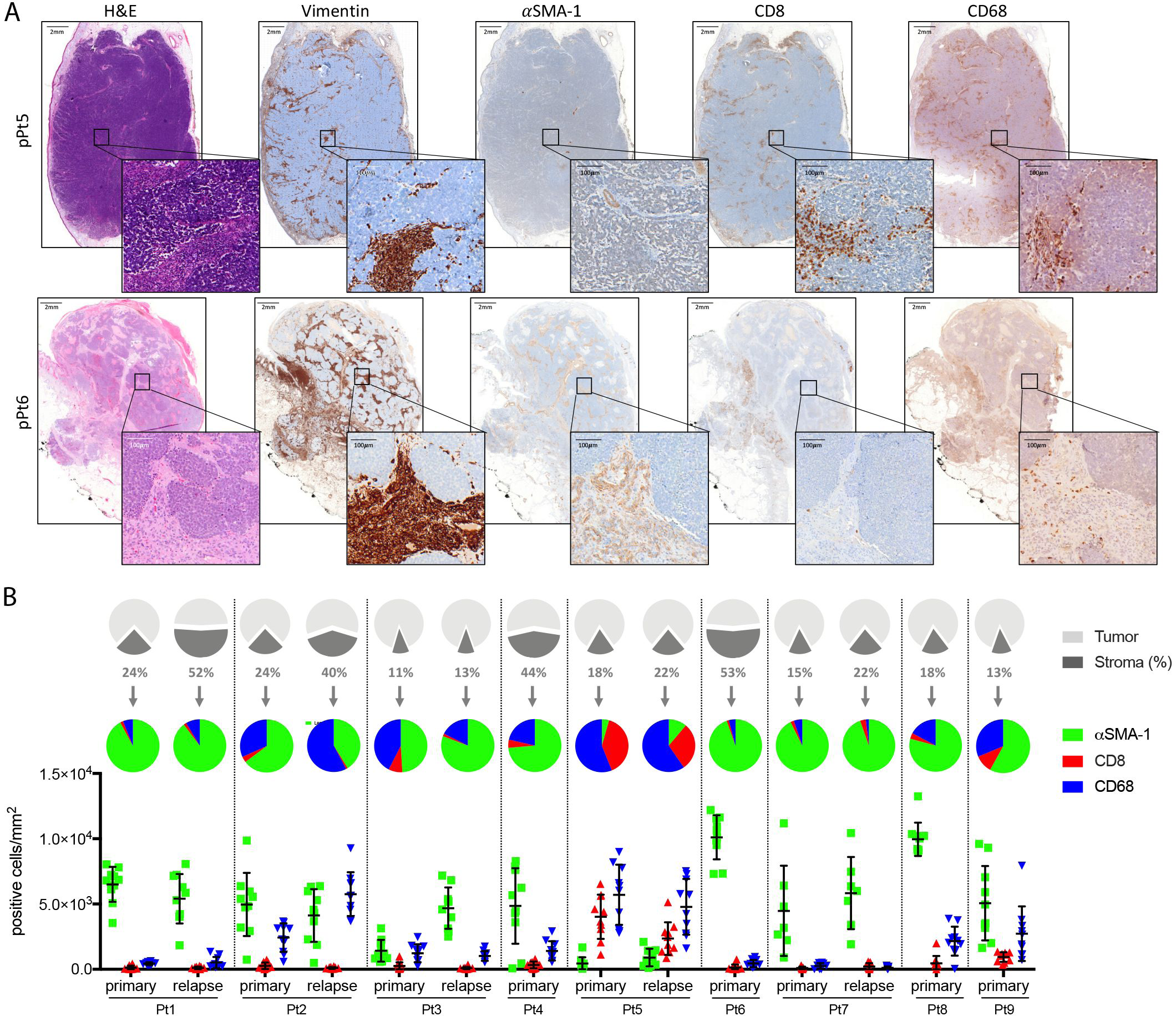
Characterization of the tumor microenvironment in MCC patients. **A** Representative images of hematoxylin and eosin (H&E) staining and immunohistochemical staining (brown) in the primary MCC from Pt5 (upper row) and Pt6 (lower row). Serial tissue sections were stained with antibodies against vimentin, αSMA-1, CD8, and CD68, respectively. Scale bars: 2 mm. Black squares insets show 25 x magnification; scale bar: 100 μm. **B** The intratumoral stromal streaks in the original MCC from the study cohort—primary and relapsed when available—were quantified with a pixel classifier trained in QuPath. The streaks are represented as pie charts with stromal components (dark grey) given as percentages (upper row). Pie chart and scatter plots showing the number of cells per mm^2^ that stained for αSMA- 1^+^ (green), CD8^+^ (red), and CD68^+^ (blue) in the intratumoral stromal streaks (lower raw). The number of cells/mm^2^ was quantified from 10 independent tumor areas (represented as individual point) using the fast cell count tool by QuPath.

### Patient-derived CAFs differentially enhance tumor growth and metastasis formation when co-injected with MKL-1 cells into SCID mice

To understand the impact of MCC-derived CAFs on tumorigenesis *in vivo*, we carried out xenograft mouse experiments by injecting CAFs together with tumor cells into an orthotopic site and directly assess the effects of CAFs on tumor growth (Albrengues *et al*, 2015). We employed an MCC xenograft metastasis mouse model that had been previously shown to develop spontaneous metastasis upon subcutaneous engraftment of the MCC cell lines, including MKL-1 (Knips *et al*, 2017). Patient-derived CAFs (Table 1) were subcutaneously inoculated into SCID mice (n= 3) at a 1:1 ratio with 1 x 10^6^ MKL-1 cells. Control experiments included the injection of MKL-1 cells alone (n=3) or at a 1:1 ratio with HFFs (n=3). With the exception of one out of three mice injected with CAFs from rPt5, which died soon after the injection for unrelated causes, mice were sacrificed at 3 months post-injection (pi) to ensure morphological and molecular characterization at a specific time point (Figure 3A). As shown in Figure 3A-C and EV2, tumors derived from co-injection of MKL-1 cells and rPt2, pPt4, or rPt5 CAFs exhibited the highest tumor growth rate and weight, while injection of MKL-1 together with rPt7 CAFs gave rise to intermediate-size tumors. However, when we co-injected MLK1 cells with CAFs derived from rPt3, nPt6, pPt8, or pPt9, we observed tumors with growth parameters similar to those obtained by injecting MKL-1 cells alone or with HFFs (Figure 3 A-C and Figure EV2).

**Figure 3.**
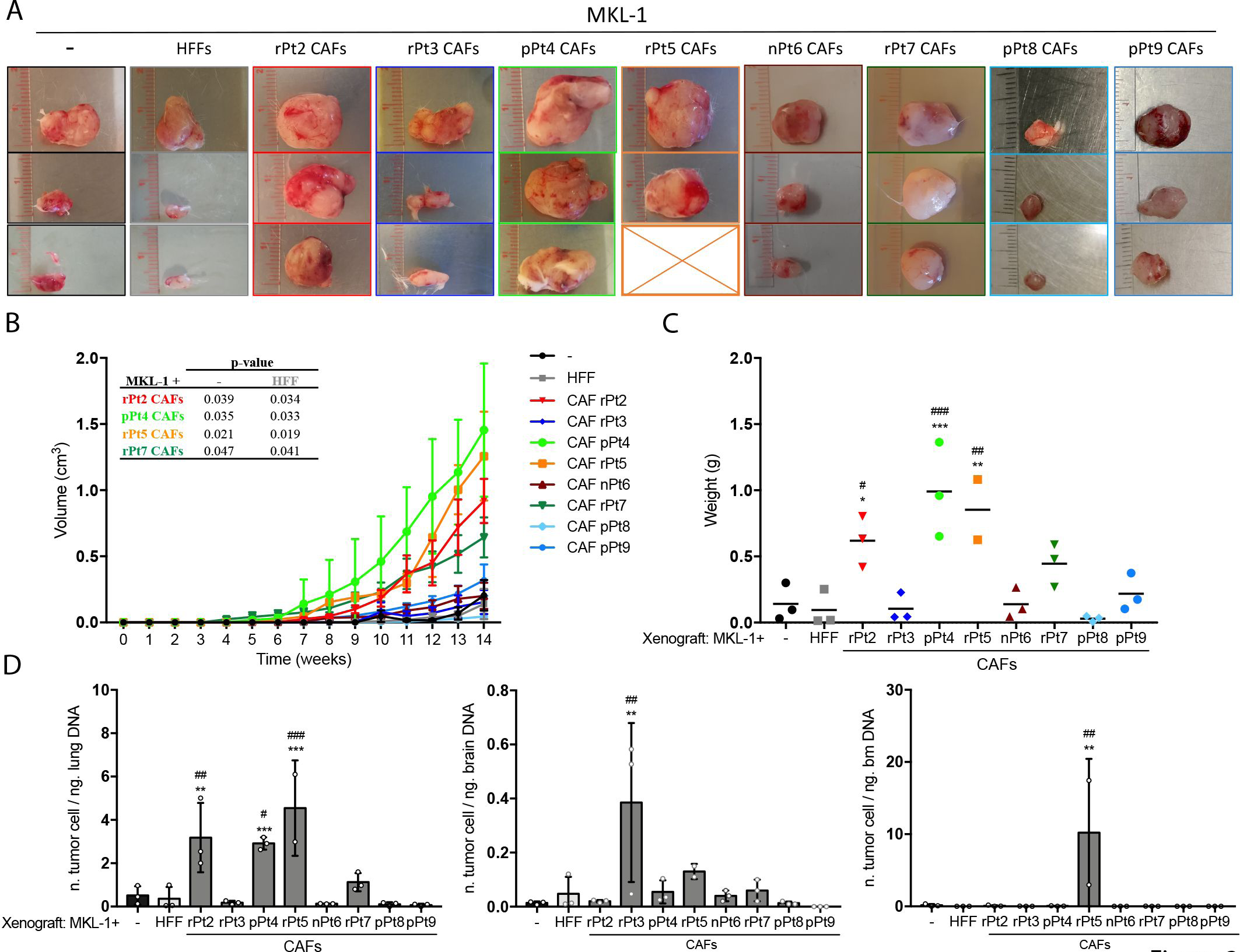
Patient-derived CAFs significantly enhance MKL-1 xenograft tumor growth and metastasis formation into SCID mice. **A** The xenograft tumors grown into SCID mice at the endpoint of the experiment (three months) are shown from three individual mice, with the exception of rPt5 CAFs (n=2), obtained by injecting MKL-1 cells alone (-) or with 10^6^ HFFs or CAFs (rPt2 to pPt9) into SCID mice. Images of the explanted xenografts are shown (*n* = 3 for each combination, with the sole exception of rPt5 *n* = 2). **B** Tumor growth was weekly assessed, and growth curves are represented as mean ± SD for each group. Statistical analysis of MKL-1/CAFs (rPt2 to pPt9) was performed by two-way ANOVA *vs* either MKL-1 alone (-) or MKL-1/HFF curves. *P*-values, when significant, are indicated in the table. **C** Tumor weight in each tumor was evaluated at 3 months after injection. **D** Quantification of metastatic cell number in lungs, brain, and bone marrow of engrafted mice as determined by real-time qPCR for human Alu sequences. The number of cancer cells was normalized to the input mouse DNA ng. Data Information: In (C, D), results are represented as mean values of the different mouse replicates; each symbol represents an individual animal. Statistical analysis was performed by classical one-way ANOVA followed by post-hoc Bonferroni correction. Significance values of MKL-1/CAFs (rPt2 to pPt9) *vs* MKL-1/HFFs: \**P* < 0.05, \*\**P* < 0.01, \*\*\**P* <0.001 or *vs* MKL-1: ^#^*P* < 0.05, ^##^ *P* < 0.01, ^###^*P* < 0.001.

We next assessed metastasis formation in different mouse organs (i.e., lung, brain, and bone marrow) by amplifying human Alu sequences, as previously published (Knips *et al*., 2017). Consistent with the tumor growth findings, the number of metastatic cells in the lungs of mice co-injected with MKL-1 cells and rPt2, pPt4, or rPt5 CAFs was significantly higher than that of mice injected with MKL-1 cells alone or in combination with HFFs or CAFs from rPt3, nPt6, rPt7, pPt8, or pPt9 (Figure 3D, left panel). In contrast, we only found few metastases in the brain and bone marrow in mice co-injected with MLK1 cells and CAFs from rPt3 or rPt5, respectively (Figure 3D middle and right panels). Of note, Pt3 died of brain metastasis three months after surgical removal of a relapsed MCC.

Overall, our results suggest that MCC-derived CAFs promote tumor progression and metastasis, although some degree of heterogeneity does exist among the patients analyzed.

### Enhanced tumor growth correlates with increased number of CAFs and angiogenesis in mouse xenografts

To further characterize the differential ability of CAFs to promote tumor growth in our mouse xenografts, we measured CAF density, CAF distribution, and the extent of angiogenesis. For this purpose, we took advantage of an antibody that specifically recognizes human vimentin in IHC, as demonstrated by the lack of signal in xenografts derived from MKL-1 cells alone (Figure 4A, upper left panel). IHC revealed significant differences in fibroblast density among xenografts. In particular, the fast-growing xenografts derived from MKL-1 with rPt2, pPt4, rPt5, or rPt7 CAFs showed significantly increased Vim^+^ staining compared with slow-growing xenografts (Figure 4A, left panels, and Figure 4B). Similarly, vessel number (Figure 4A, middle panels, and Figure 4C) and diameter (Figure 4D) were increased as shown by immunostaining for CD31. Serial sections were then co-stained by immunofluorescence with two antibodies, one specific for human Vim (green) and both, human and mouse Vim (red). This co-staining indicated that human fibroblasts (green) were accumulated in the fast-growing xenografts around the large blood vessels, in close contact with the murine endothelial cells of the vessel walls (red). Quantification of the number of vessels exhibiting this colocalization revealed that xenografts injected with MKL-1 cells plus rPt2, pPt4, rPt5, or rPt7 CAFs showed a high percentage of vessels surrounded by large (black) or medium (dark gray) numbers of human fibroblasts. By contrast, xenografts injected with control cells or with MKL-1 plus pPt8 or pPt9 CAFs mainly displayed vessels surrounded by only a small number of human fibroblasts (light gray, Figure 4E). Our data point to a positive correlation between the percentage of human fibroblasts surrounding the vessels and the vessel diameter (Figure 4F).

**Figure 4.**
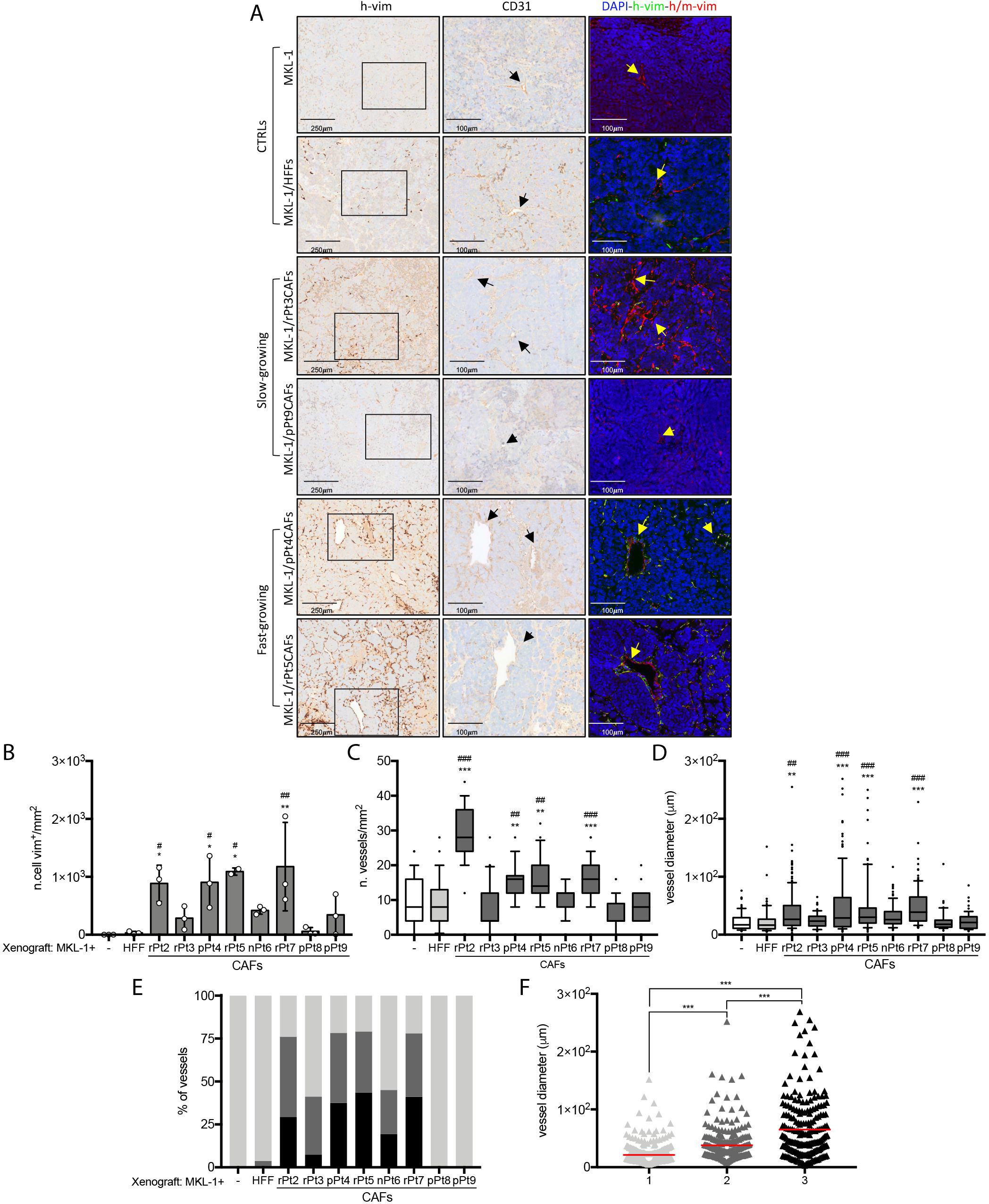
MCC-derived CAFs potentiate angiogenesis in MKL-1 xenografts. **A** Representative images of serial tissue sections from FFPE blocks of the xenografts obtained by subcutaneous injection of MKL-1 cells alone (-), MKL-1 plus normal fibroblasts (HFFs), or MKL-1 plus rPt3, pPt9, pPt4, or rPt5 CAFs. Tissue samples were stained with antibodies against human vimentin (left panels) or CD31 (central column). The right panels show the immunofluorescence analysis of serial sections co-stained with anti-vimentin antibodies as follows: human-specific (h-vim, green) and anti-human/mouse (h/m-vim, red). The nuclei were counterstained with DAPI (blue). The area shown in the central and right panels is a magnification of the area contained in the respective hollow rectangle. Vessels are highlighted with arrows. **B** The number of h-vim^+^ cells/mm^2^ was measured in the entire tumor mass sections of all xenografts using the fast cell count tool from QuPath software. **C** Vessel density in sections stained with antibodies against CD31. Microvessels were counted manually in at least 8-10 fields per tumor section at ×200 magnification. **D** Vessel size was determined by measuring the wider vessel diameter with QuPath software. **E** Correlation of vessel diameter with number of CAFs found in close proximity to vessels. Vessel quantification was performed as described in (D) and a score of 1 (low density, light gray), 2 (medium density, dark gray), and 3 (high density, black) was given based on the density of h- vim^+^ cells surrounding the vessels. The number of blood vessels, scored 1 to 3 in each explanted tumor, is represented as the average percentage within the replicates (*n* = 3 for each combination, with the sole exception of MKL-1/rPt5CAFs, *n* = 2). **F** For each blood vessel, regardless of the mouse group, the size (y-axes) is plotted against its corresponding score (x-axes). Data Information: in (B) data are represented as mean ± SD, and each dot represents an individual animal; In (C and D), the whisker-box plots represent the 25^th^-75^th^ percentiles, with midlines indicating the median values, whiskers extending to the 10^th^-90^th^ percentiles, and dots showing outliers. Significant levels are \**P* < 0.05, \*\**P* < 0.01, \*\*\**P* < 0.001 *vs* MKL-1/HFFs or ^#^*P* < 0.05, ^##^ *P* < 0.01, ^###^*P* < 0.001 *vs* MKL-1 in (B, C, and D) and \*\*\**P* < 0.001 in (F) from classical one-way ANOVA followed by post-hoc Bonferroni correction.

### The transcriptome profile of the xenografts derived from growth-promoting CAFs is characterized by a pro-angiogenic signature

In order to unravel CAF-mediated functions in promoting tumor formation, we performed bulk transcriptome sequencing (RNA-Seq) of xenografts obtained by injecting MKL-1 cells with CAFs from rPt2, pPt4, or rPt5, which we had previously shown to promote enhanced tumor formation (Figure 3). As negative controls (CTRLs), we injected mice with MKL-1 cells alone or with HFFs, which were treated as biological duplicates based on principle component analysis (Appendix Figure S1). Differential gene expression analysis of CAFs *vs* CTRLs revealed 234 differentially expressed genes (DEGs), of which 98% were upregulated (*n* = 227) and only 2% downregulated (*n* = 3; Figure 5A-B) [false discovery rate (FDR) < 0.05; Log2FC > 1]. Functional annotation of DEGs (Figure 5C and Appendix Table S1) showed significant enrichment for angiogenesis-related biological processes, such as blood vessel development and morphogenesis and vasculature development. Furthermore, gene set enrichment analyses (GSEA) of the DEGs involved in the top 10 annotated gene ontology (GO) terms (Figure 5C and Appendix Table S1) showed a positive normalized enrichment score (NES) and a significant FDR < 0.25 for all the gene set lists analyzed, thereby confirming their distribution in the top portion (upregulated) of the gene dataset (Appendix Figure S2). Biological pathway enrichment analysis was also performed through KEGG pathway analysis with DEGs being highly associated with pathways defined as PI3K-Akt signaling pathway and ECM-receptor interaction (Figure 5D), which have been previously linked to MCC carcinogenesis and CAF functions in tumorigenesis (Hafner *et al*, 2012; Kalluri, 2016; Nardi *et al*, 2012).

**Figure 5.**
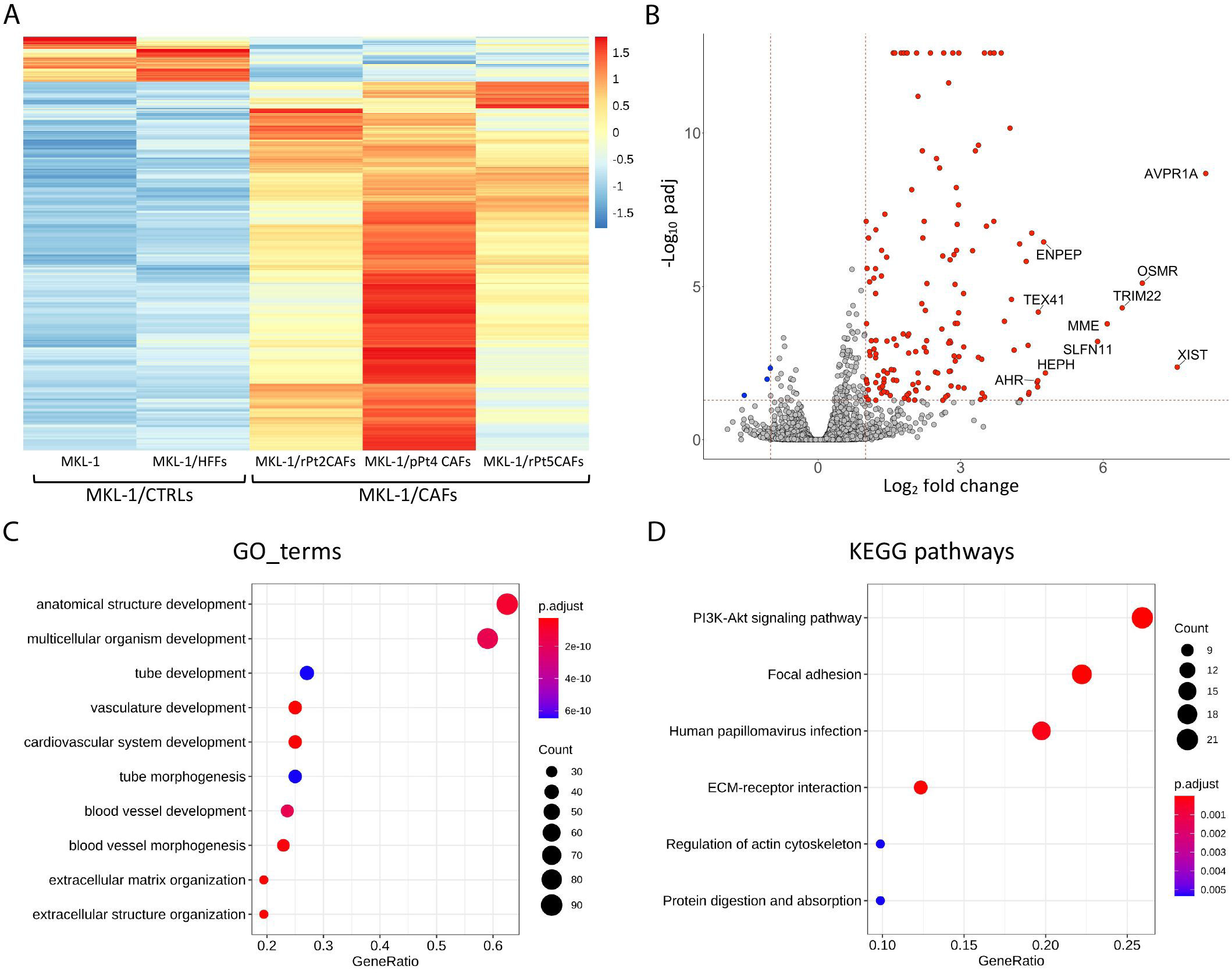
Differential gene expression profiling of xenografts injected with MKL-1/CAFs from rPt2, pPt4, and 5 *vs* those injected with MKL1/HFFs or MKL-1 alone. **A** Heatmap of RNA-Seq data showing genes resulting differentially expressed (log2FC |>1, *P* adj < 0.01) between tumor CAF (MKL-1/CAFs from rPt2, pPt4, and rPt5) and CTRL (MKL-1 and MKL-1/HFFs) groups. **B** Volcano plot of RNA-Seq data showing up- and downregulated genes in MKL-1/CAFs *vs* MKL-1/CTRLs. Significantly deregulated genes (DEG; log2FC |>1, *P* adj < 0.01) are visualized as red dots. The vertical bars delimitate log2FC |1, while the horizontal ones delimitate -log10 padj = 1.3. **C** GO term analysis. DEG genes resulting from CAFs *vs* CTRLs were analyzed using the DAVID bioinformatics resources. Circle size maps the number of genes in each GOterm/KEGG pathway, whereas colors map significance (*P* adjust). **D** KEGG pathway enrichment analysis. This analysis was performed as described above.

To broaden our analysis, we investigated the DEGs using ingenuity pathway analysis (IPA). IPA analysis predicted a wide range of activated and decreased disease and function in CAFs compared to CTRLs (Figure EV3 and Appendix Table S2). More specifically, 68 disease and function annotations were defined as being activated (Figure EV3, orange bars), including growth, migration, and invasion of tumor cells, development of vasculature, and angiogenesis, while 16 annotations displayed a decreased activation state (blue bars), including necrosis, apoptosis, and cell death of carcinoma cell lines.

**Table 2.**
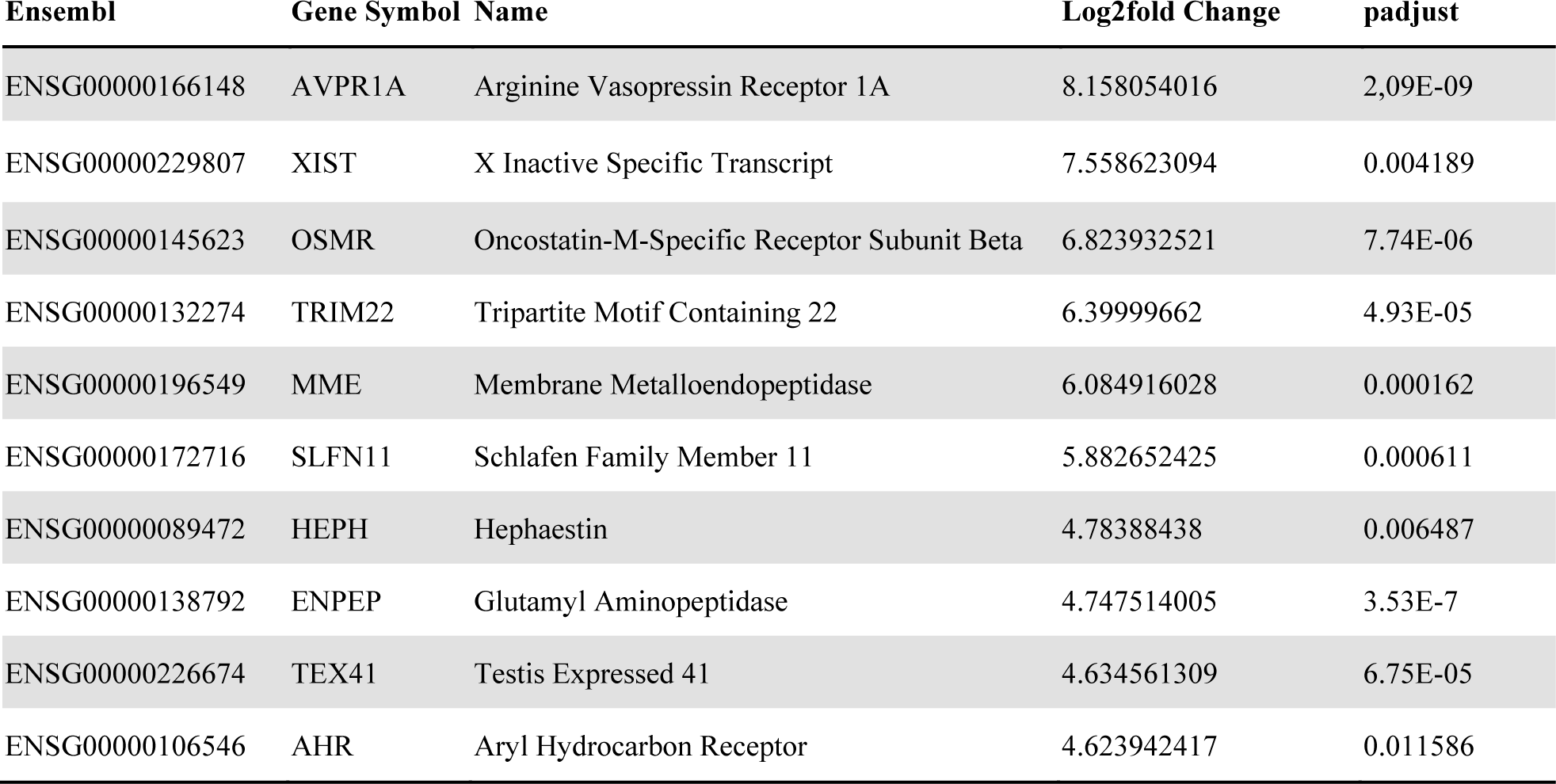
Top 10 overexpressed genes in xenograft tumors derived from mice engrafted with MKL-1/rPt2, pPt4, or rPt5 CAFs compared to controls (MKL-1 and MKL-1/HFFs)

Overall, the transcriptome results from the xenografts injected with tumor-promoting CAFs support the involvement of CAFs in tumor development and progression and, in particular, suggest their engagement in angiogenesis processes in MCC.

### CAF-induced angiogenesis goes through the APA/AngII-III/AT_1_R pathway

To determine the functional role of the pro-angiogenic signature revealed by the transcriptome analysis, we first assessed the expression levels of the top 10 upregulated genes (Table 2) by RT-qPCR. In particular, we focused our attention on the arginine vasopressin receptor 1A (*AVPR1A*) — here the most upregulated one— previously shown to play a pivotal role in prostate cancer progression and metastasis formation (Zhao *et al*, 2019), and on the glutamyl aminopeptidase (*ENPEP*) and membrane metalloendopeptidase (*MME*) genes, which were among the GO term angiogenesis genes listed in Appendix Table S1. Quantitative RT-PCR analysis, performed using total RNA re-extracted from all the xenografts—with the exclusion of pPt4 and pPt8 CAF-injected xenografts due to insufficient RNA quality—, revealed that all three genes were upregulated, albeit to different extents, in the xenografts co-injected with MKL-1 and CAFs compared to MKL-1 alone or MKL-1 plus HFFs (Figure 6A). Importantly, their expression levels were significantly higher in MKL-1/rPt2, rPt5, or rPt7 CAF tumors than in MKL-1/rPt3, nPt6, or pPt9 tumors, suggesting acorrelation between upregulation of *AVPR1A, ENPEP* and *MME* and increased tumor burden. Of note, *AVPR1A, ENPEP* and *MME* expression levels also correlated with the number of Vim^+^ cells in the xenografts (Figure 6B). While in MKL-1 cells *ENPEP*, *MME,* and *AVPR1A* were all transcriptionally inactive, *ENPEP* and *MME* were expressed in HFFs and patient-derived CAFs (Figure 6C). Interestingly, *AVPR1A* gene expression was only detected in xenografts (Figure 6A) but not in the 2D cell cultures of all the cell lines and primary cells analyzed.

**Figure 6.**
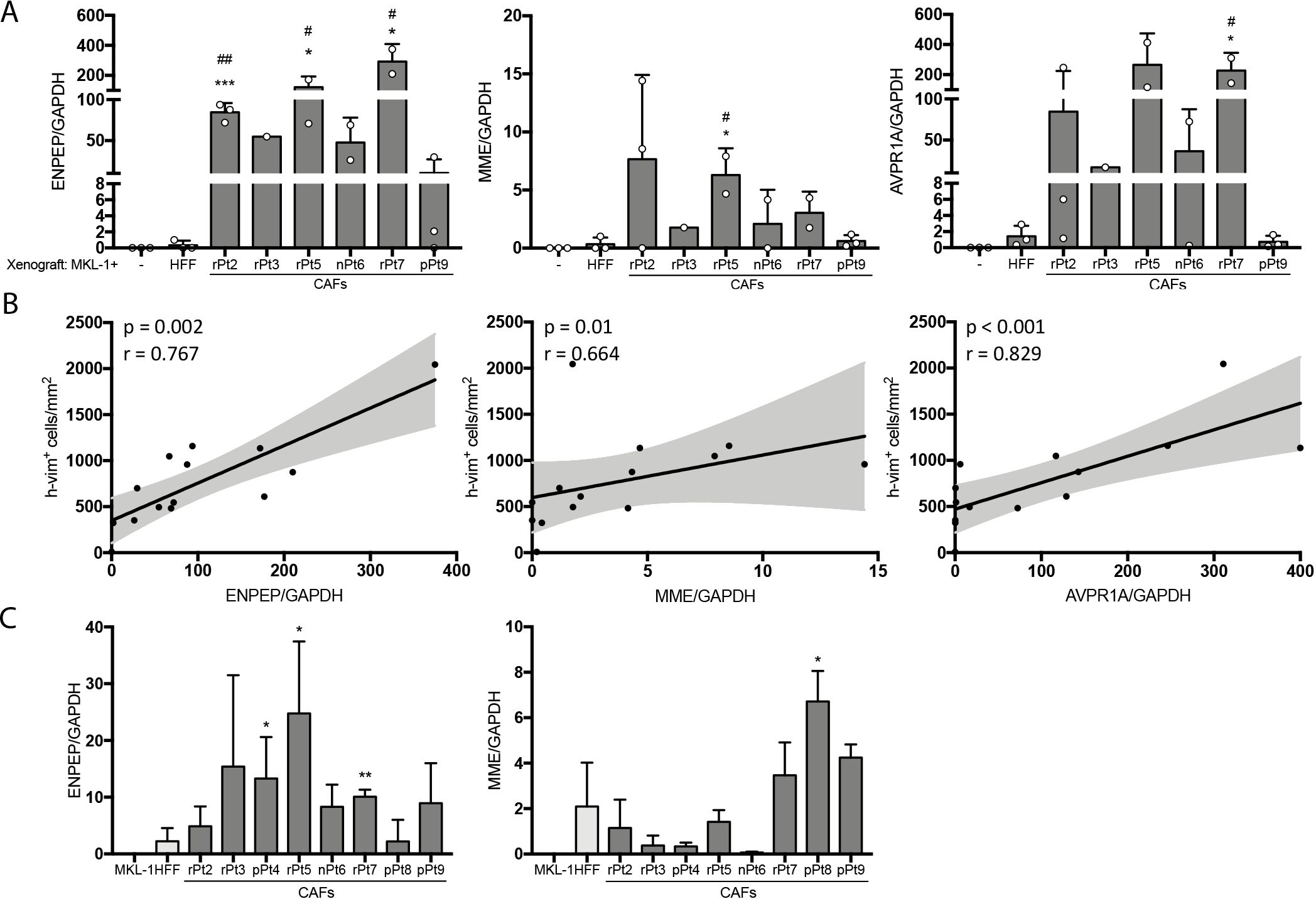
AVPR1A, ENPEP and MME is differentially expressed by fast-growing xenografts. **A** qRT-PCR analysis for ENPEP, MEE and AVPR1A mRNA expression levels in total RNA extracts from individual explanted xenografts. Values were normalized to GAPDH and plotted as fold induction over MKL-1 plus HFF-injected tumors. Data are represented as means ± SD, and each dot represents an individual animal. Statistical analysis was performed by one-way ANOVA followed by post-hoc Bonferroni correction. Significant levels: \**P* < 0.05, \*\*\**P* < 0.001 *vs* MKL-1/HFFs or ^#^*P* < 0.05, ^##^*P* < 0.01 *vs* MKL-1. **B** Spearman rank correlation of h-vim^+^cell/mm^2^ (y-axes) with ENPEP, MEE and AVPR1A mRNA expression levels (x-axes). p (*P*-value) and r (spearman correlation coefficient) are indicated in the graph. **C** qRT-PCR analysis for ENPEP and MME mRNA expression levels from CAFs, HFFs, or MKL-1. Values were normalized to GAPDH and plotted as fold induction over HFFs. Data are represented as means ± SD. Statistical analysis was performed by one-way ANOVA followed by post-hoc Bonferroni correction. Significant levels: \**P* < 0.05, \*\**P* < 0.01 *vs* HFFs.

*ENPEP* encodes the membrane aminopeptidase A (APA), a membrane-bound metallopeptidase that converts bioactive peptides such as angiotensin II (Ang II) into angiotensin III (Ang III) (George *et al*, 2010). Both peptides are known to exert pro-angiogenic activity *via* the angiotensin II type 1 receptor (AT_1_R) (Bosnyak *et al*, 2011; George *et al*., 2010; Reaux *et al*, 2001). In addition, Ang II conversion into Ang III prevents Ang II cleavage into Ang(1-7) by ACE2, which in turn binds the MAS receptor, promoting an anti-angiogenic effect (Catarata *et al*, 2020). Thus, we decided to test whether pharmacological inhibition of this pathway would impair CAF-induced angiogenesis in an *in vitro* model of tubulogenesis. Human dermal blood endothelial cells (HDBECs) were grown on a monolayer of patient-derived CAFs or HFFs, and their ability to form tubule-like structures was assessed after 10 days. All patient-derived CAFs supported *in vitro* tube formation, whereas non-activated HFFs failed to generate tubular structures (Figure EV4). For all subsequent *in vitro* tube formation assays, we used rPt-5 CAFs due to their robust tumor growth-promoting and angiogenic properties. The APA inhibitor amastatin was used at concentrations of 1, 10, and 100 μM. Lack of cell toxicity was assessed by MTT assay (Appendix Figure S3). As shown in Figure 7 and Figure EV5, when amastatin was added to the co-cultures, we observed a dose-dependent inhibition of tubule-like structure formation, measured as mean tubule length, total tubule length, and tubule density upon CD31 staining (Figure 7A-B). Similarly, tube formation was inhibited when the co-cultures were treated with the AT_1_R inhibitor candesartan (Figure 7C-D and Figure EV5) (Uemura *et al*, 2008). We did not observe a synergistic effect when co-cultures were treated simultaneously with the two inhibitors (Figure EV5).

**Figure 7.**
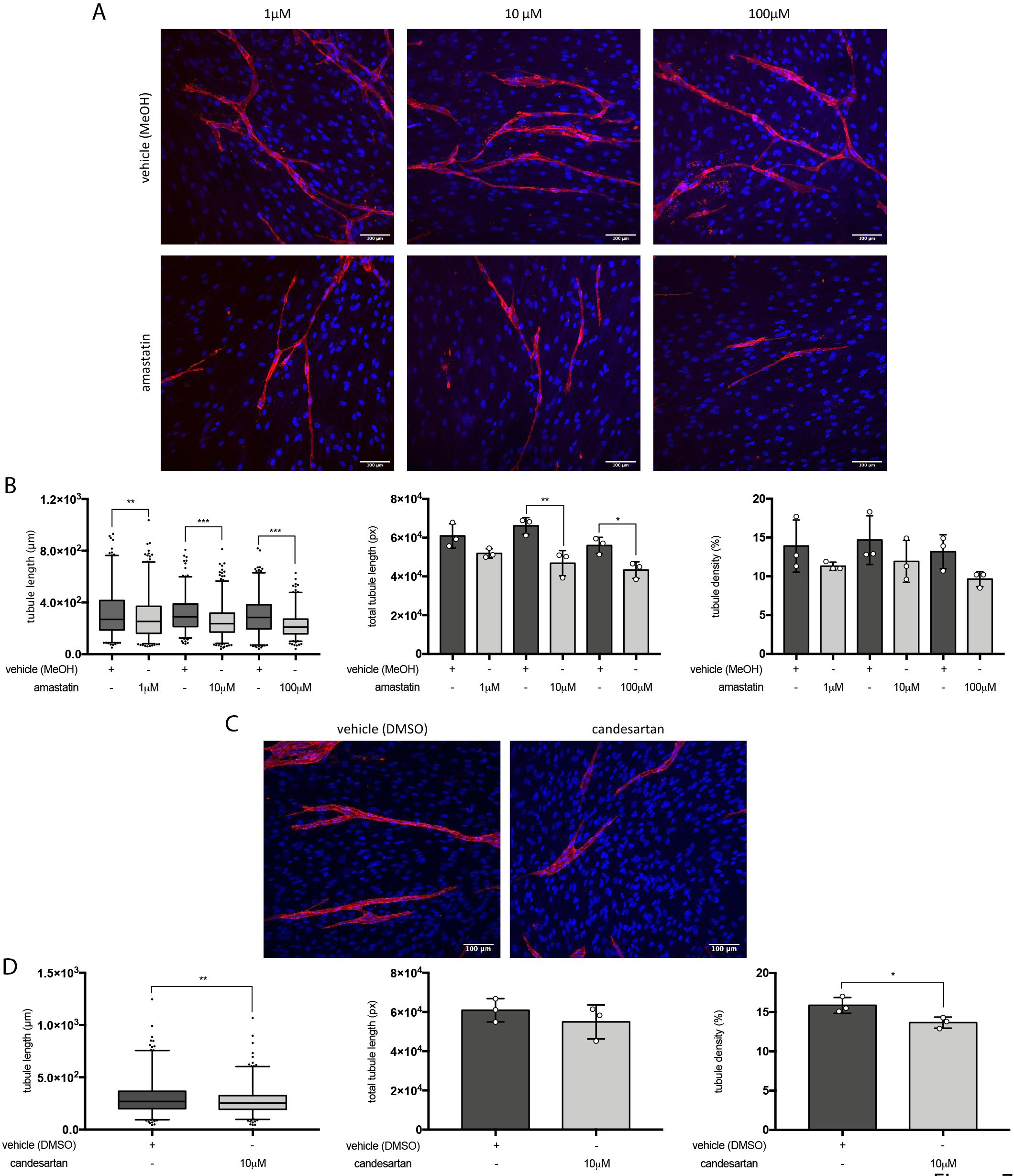
Pharmacological inhibition of the APA/AT_1_R pathway impairs CAF-induced tubulogenesis. **A** Representative confocal images of HDBECs co-cultured with rPt5 CAFs for 10 days in the presence of amastatin or vehicle as control (MeOH) at the indicated concentrations. Scale bar = 100 μm. **B** Quantification of mean tubule length (left panel), total tubule length (central panel), or tubule density (right panel) from the co-cultures shown in panel A. Data represent mean ± SD of three independent experiments, one-way ANOVA followed by post-hoc Bonferroni correction. Significant levels: \**P* < 0.05, \*\**P* < 0.01, \*\*\**P* < 0.001. **C** Representative confocal images of HDBECs co-cultured with rPt5 CAFs for 10 days in the presence of candesartan (10 μM) or vehicle as control (DMSO). Scale bar = 100 μm. **D** Quantification of mean tubule length (left panel), total tubule length (central panel), or tubule density (right panel) from the co-cultures shown in panel C. Data represent mean ± SD of three independent experiments, unpaired *t* test; \**P* < 0.05, \*\**P* < 0.01.

Altogether, these findings clearly indicate that the APA/AT_1_R pathway is involved in the pro-angiogenic activity of MCC-derived CAFs.

## Discussion

There is growing evidence that cancer progression is supported by changes in the tumor microenvironment, where CAFs are the most abundant cell type (Kalluri, 2016). While the protumorigenic role of CAFs in various solid tumors has been clearly established, the function of these cells in MCC initiation and progression remains poorly understood. The present study describes a novel mechanism of MCC etiopathogenesis based on CAF-driven tumor growth and angiogenesis *via* the APA/Ang II-III/AT_1_R axis, in which APA expression in CAFs potentially accounts for this major contribution.

Our molecular analysis shows that MCC-derived CAFs are characterized by increased αSMA-1 expression as well as enhanced contractility and secretion of pro-inflammatory cytokines, hallmarks of an activated phenotype. Furthermore, 6 out of the 9 MCCs analyzed in our study were positive for MCPyV, which together with UV light is the main etiologic factor for MCC (DeCaprio, 2021). Although we did not find any specific correlations between CAF activity and/or phenotype and MCPyV positivity, most of these MCC-derived CAFs, in contrast to non-activated fibroblasts, significantly enhanced tumor growth of MKL-1 cells in immunosuppressed mice. Notably, xenografts generated by co-injection of rPt2, pPt4, rPt5, or rPt7 CAFs and MKL-1 cells revealed increased tumor volume and weight compared with xenografts generated with rPt3, nPt6, pPt8, or pPt9 CAFs plus MLK-1 cells. Consistent with an active role of CAFs in enhancing tumor progression, an increased number of lung metastases were observed in mice injected with rPt2, pPt4, or rPt5 CAFs plus MKL-1 cells. Another interesting finding is that we could only detect brain metastases in mice injected with CAFs derived from Pt3, who had died from brain metastasis three months after the surgical resection of the tumor. Although this was only one case, the apparent correlation between CAF-driven brain metastasis and poor disease outcome further strengthens the impact of CAFs on MCC progression.

An important consideration from our *in vivo* tumorigenicity assays is that patient-derived CAFs may differentially affect MCC growth and metastasis, implying that their *in vivo* acquired activated phenotype along with their intrinsic heterogeneity in driving tumor growth and metastasis are fully preserved in our culture conditions. More specifically, the observed enhanced xenograft growth correlated with increased stromal cells density in the xenografts, as documented by the quantification of Vim^+^ cells in tissue sections. Of note, we found that Vim^+^ cells were more abundant in fast-growing than in slow-growing xenografts, and that in the latter co-injection of CAFs did not significantly increase tumor growth induced by MKL-1 cells alone. Consistent with this observation, the number of human fibroblasts detectable in the xenografts where normal fibroblasts were co-injected with MKL-1 cells was extremely low. The observed increased CAF density was accompanied by augmented vessel density and size, suggesting that CAFs may actively contribute to vascularization, favoring tumor growth. Another interesting observation supporting the pro-angiogenic function of CAFs is that the areas near the blood vessels in the rapidly growing xenografts were densely populated by human fibroblasts.

Comparative gene expression profiling by whole genome RNA-sequencing of xenografts injected with the growth-promoting rPt2, pPt4, or rPt5 CAFs *vs* xenografts injected with HFFs or MKL-1 cells alone showed significant enrichment in angiogenesis-related biological processes, such as blood vessel development and morphogenesis and vasculature development. Among the top ten most upregulated genes, the mRNA levels of *AVPR1A*, *ENPEP* and *MME* were significantly increased in fast-growing *vs* slow-growing xenografts. The fact that *MME* and *ENPEP* expression was increased in CAFs *vs* HFFs and totally absent in MKL-1 cells cultured as monolayers strongly suggests that CAFs are the major site of their expression. Intriguingly, while *AVPR1A* gene expression was confirmed at the RNA level in both xenografts (Figure 5E) and patient-derived MCC specimens (Appendix Figure S4), we failed to detect its expression in both HFFs/CAFs and MKL-1 cells grown *in vitro* as either monolayer or co-cultures. Thus, further experiments *in vivo* are clearly needed to ascertain whether AVPR1A is induced in CAFs or tumor cells and plays a role in MCC progression and metastasis.

*ENPEP* encodes APA, a homodimeric type II membrane-spanning cell surface protein with a zinc metallopeptidase activity that hydrolyzes N-terminal glutamyl or aspartyl residues from oligopeptide substrates (Nanus *et al*, 1993). The best understood role of APA is the conversion of angiotensin II (Ang II) to angiotensin III (Ang III) in the renin-angiotensin system (RAS) (Nanus *et al*., 1993). Most Ang II and Ang III actions involve AT_1_R, a G protein-coupled receptor, which is upregulated in many cancer types (Bosnyak *et al*., 2011; George *et al*., 2010). In the brain, experiments involving APA and APN specific inhibitors have provided clear evidence that Ang III is the main effector peptide involved in AT_1_R activation and vasopressin release (Zini *et al*, 1996). In addition, the blockage of Ang II conversion to Ang III with APA inhibitors has been shown to favor the activation of other metabolic pathways, such as that involving the angiotensin-converting enzyme 2 (ACE2) and the Mas receptor ligand, whose activity is known to inhibit cancer growth and angiogenesis (Reaux *et al*., 2001; Wright *et al*, 2012).

Although RAS was originally discovered for its relevant role in regulating vascular homeostasis, emerging data from *in vitro*, animal, and clinical studies indicate that aberrant regulation of this pathway can promote tumor growth and angiogenesis in several tumor types (George *et al*., 2010). Conversely, RAS inhibition can suppress tumor growth, metastasis, and angiogenesis in a variety of tumor models (Pinter & Jain, 2017). Fittingly, APA is upregulated in the perivascular cells of angiogenic tumor vessels where it promotes angiogenesis, while in normal blood vessels it is only barely detectable (Marchio *et al*, 2004).

In this study, we show that the *ENPEP* gene encoding APA is transcriptionally upregulated in fast-growing xenografts where it plays a crucial role in CAF-induced angiogenesis. We also demonstrate that APA inhibition with the antagonist amastatin impairs tubule formation. This inhibitory effect is dose-dependent and primarily affects the length of tubule-like structures as well as their extension and density rate, indicating that the tubulogenesis activated by the CAF co-culture strictly relies on APA activity. Given that the two enzymatic byproducts of APA, Ang II and Ang III, mainly function through AT_1_R, it is not surprising that treatment of CAF/HDBEC co-cultures with the AT_1_R inhibitor candesartan impaired the formation of branched, interconnecting networks of endothelial cell tubules, as we have demonstrated in amastatin-treated cultures.

Given that very few studies have thoroughly addressed the biological significance of stromal cells in MCC progression, our investigation is the first to provide evidence that these cells may play a relevant role in modulating tumor growth, particularly angiogenesis. This new paradigm would be consistent with a recent report showing that intercellular communication between MCC cells and stromal fibroblasts is required for the generation of activated/polarized CAFs (Fan *et al*., 2021). Consistent with our findings, variability/heterogeneity of stromal components populating MCC lesions has been reported, a phenomenon also observed in many other cancers. This aspect is particularly important considering that, to date, no reliable prognostic markers for MCC have been identified. Thus, it is tempting to speculate that quantification/characterization of the stromal components in MCC may be used to predict more accurately disease progression.

In conclusion, our study highlights the biological role of CAFs in MCC development and metastasis formation. The growth-promoting activity of MCC-derived CAFs is mostly due to increased angiogenesis. Mechanistically, we demonstrate that this process is largely mediated by the APA/Ang II-III/AT_1_R axis, with APA expression in CAFs being the upstream triggering event, suggesting a potential role of APA as a prognostic and therapeutic marker for MCC. Further experiments will be necessary to determine the effect of APA inhibitors, alone or in combination with chemotherapy, on tumor growth, angiogenesis, and metastasis formation *in vivo*.

## Materials and Methods

### Patients and cell culture

Nine patients with a histological diagnosis of Merkel cell carcinoma (MCC) who underwent surgical excision of primary or relapsed tumor or lymph nodal metastasis were included in this study. They were recruited between July 2015 and April 2019 through “Rete Oncologica Piemontese” from various Hospitals in the Piedmont region (Northern Italy). All patients were informed at the time of the surgery that their clinical data and specimens (i.e., FFPE blocks and fresh tissue) would be used for research purposes and signed a written informed consent form. Study approval was obtained from the Ethics Committee of “Maggiore della Carità” Hospital (agreement number 213/CE). Fresh human MCC specimens were obtained following surgical excision of three primary tumors, four relapsed tumors, and one lymph node metastasis.

CAFs were directly isolated from the patient tissue as previously described (Lau *et al*, 2016). Briefly, tumors were washed twice with PBS, visible excess of fat tissue was removed, and tissues were finely minced using a sterile scalpel. Homogenized tissues were incubated in RPMI media supplemented with 10% FBS and P/S and maintained at 37°C in 5% CO_2_. After 24 h, the unattached cells were removed, and the remaining cells were allowed to grow on the plate for 2- 3 weeks with the media replenished every other day. When adherent cells reached confluence, they were trypsinized and transferred into a new culture flask and grown in DMEM supplemented with 10% FBS and P/S. For functional assay, only low passage numbers were used.

The MCPyV-positive MCC cell line MKL-1 (Rosen *et al*, 1987) was purchased from Sigma-Aldrich (Cat. No. 09111801) and grown in RPMI 1640 (Gibco) supplemented with 10% fetal bovine serum (FBS, Sigma-Aldrich, Milan, Italy), 100 U/ml penicillin, and 0.1 mg/ml streptomycin (P/S). HFFs (ATCC Cat. No. SCRC-1041) were maintained in DMEM 10% FBS and P/S. HDBECs (PromoCell, Cat. No. C-12211) were grown in endothelial cell basal medium (PromoCell, Heidelberg, Germany) supplemented with Endothelial Cell GM MV 2 Supplement Pack (PromoCell, Heidelberg, Germany).

### Collagen contractility assay

Collagen contractility assay was performed as previously described (Saraswati *et al*, 2019). Briefly, 3 x 10^5^ HFFs or CAFs were embedded in 750 μl of rat tail collagen-I (Corning, Germany) supplemented with 1.5 ml of serum-free DMEM and 12.5 μl of 1N NaOH. Six-hundred μl of cell/collagen mix was plated in 24-well plates and incubated for 15-20 min until gelled. The gel was then separated from the walls of the well with the help of a 30G needle, and 600 μl/ well of DMEM supplemented with 10% FBS was added on top of the gel. Gels were incubated for 24 h at 37°C, fixed with 4% PFA, and stained with eosin. Well and gel diameters were measured using Fiji (Schindelin *et al*, 2012). The percentage of gel contraction was calculated using the formula 100 x (well diameter - gel diameter)/well diameter.

### Immunofluorescence, immunohistochemistry, and immunoblotting

Consecutive 5-μm thick tissue sections were obtained from FFPE blocks. Hematoxylin and eosin (H&E) and immunohistochemistry staining were performed using the automated immunostainer BenchMark ULTRA Stainer (Ventana Medical System, Tucson, AZ). For immunofluorescence, antigen unmasking was performed by heating the slides in a conventional decloaking chamber at 750W for 15 min, followed by an additional step at 350W for 10 min in 10 mM citrate buffer at pH 6.0 (Vector Laboratories, Burlingame, CA, USA). Primary antibodies were diluted together in 5% normal goat serum/PBS and incubated ON at 4°C. Tissue images were acquired using Panoramic Midi Scanner (3D-Hystec) and staining was quantified using QuPath software (Bankhead *et al*, 2017).

For immunoblotting, whole-cell protein lysates were prepared and processed as previously described (Gugliesi *et al*, 2005). Anti-β-actin antibodies (Mab) (Active Motif) were used as control for protein loading. Immunocomplexes were detected using sheep anti-mouse or donkey anti-rabbit immunoglobulin secondary antibodies conjugated to horseradish peroxidase (HRP) (GE Healthcare Europe GmbH) and visualized by enhanced chemiluminescence (Super Signal West Pico; Thermo Fisher Scientific) according to the manufacturer’s instructions. Images were acquired using Quantity One software (version 4.6.9; Bio-Rad Laboratories Srl). The antibodies used in this study are listed in Appendix Table S3.

### Quantitative nucleic acid analysis

MCC tumor total DNA was isolated with a QIAamp DNA Mini Kit (QIAGEN). To determine MCPyV viral load in MCC specimens, quantitative PCR (qPCR) was performed using 500 nM primers and SsoAdvanced Universal SYBR Green Supermix (Bio-Rad). The reaction conditions consisted of a 30-s 95°C enzyme-activation cycle, 40 cycles (10 s) of denaturation at 95° C, and 10 s of annealing at 60°C. Copy number analysis was completed by comparing the unknown samples with standard curves of linearized MCPyV DNA. The GAPDH DNA copy number was used as endogenous control.

Total RNA was extracted from CAFs, xenografts, and MCCs using TRIzol (Thermo Fisher Scientific, Inc.), and 1 μg per sample was retrotranscribed using iScript cDNA synthesis kit (Bio-Rad Laboratories Srl). To detect cellular gene expression, reverse-transcribed cDNAs were amplified in duplicate using SensiFast SYBR (Bioline). Real-time quantitative reverse transcription (qRT)-PCR analysis was performed on a CFX96 real-time system (Bio-Rad Laboratories Srl) under the following conditions: 95 °C for 10 min, followed by 40 cycles at 95 °C for 15 s, and 60 °C for 1 min. Relative quantification [comparative Ct (ΔΔCt) method] was used to compare the expression level of the tested genes with the housekeeping gene glyceraldehyde-3-phosphate dehydrogenase (GADPH).

To detect disseminated tumor cells in mouse organs, total DNA was extracted from the organs of the engrafted mice using QIAamp DNA Mini Kit (Qiagen), and 40 ng were amplified using human specific Alu sequences as previously described (Knips *et al*., 2017; Nehmann *et al*, 2010). Amplification was performed with a CFX96 real-time system (Bio-Rad Laboratories Srl) under the following conditions: 10 min at 95C, followed by 50 cycles at 95 °C for 5 s, 67 °C for 5s, and 72 °C for 20s. Quantification of human metastatic cells in mouse organs was determined by using standard curves generated with MKL-1 DNA dilutions ranging from 1 to 10^6^ cells/ml. The number of metastatic cells was normalized to the ng of DNA used for the amplification.

The QX200 Droplet Digital PCR System (Bio-Rad Laboratories) was used for absolute quantification of mRNA expression in MCC tumor. For each assay, a reaction mixture of 22 μL was prepared with 11 μL EvaGreen SuperMIX (Bio-Rad Laboratories), 1.1 μL of 10 μM of each primer, and 5 ng of cDNA. For droplet generation, 20 μL reaction mix was used, and the droplets were transferred to a 96-well plate. Samples were amplified in a C1000 Touch Thermal Cycler (Bio-Rad Laboratories) according to the following protocol: 95 °C for 5 s, 40 cycles of 95 °C for 30 s and 58 °C for 60 s, followed by 4°C for 5 s, 90 °C for 10 min, and 4°C for 40 s. Data were analyzed using QuantaSoft (Bio-Rad Laboratories). PCR primer sequences are detailed in Appendix Table S4.

### *In vivo* tumorigenic assay

Female SCID mice (CB17/lcr-Prkdc scid/lcrlcoCrl; Charles River Laboratories, Sulzfeld, Germany) were housed under pathogen-free conditions in our animal facilities in accordance with “The Guide for the Care and Use of Laboratory Animals”, and the experimentation was approved by the Italian Ministry of Health (Agreement number 231/2020-PR). HFFs or CAFs from Pt2 to 9 were injected along with MKL-1 cells as previously described (Knips *et al*., 2017; Lau *et al*., 2016). Briefly, 50 μl of matrigel (Corning) suspension containing 1 x 10^6^ (MKL-1 alone) or 2 x 10^6^ cells (1x10^6^ MKL-1 combined with 1x10^6^ HFFs or CAFs) were injected subcutaneously into 6-week-old SCID mice. All mice were observed and examined weekly, and tumor volumes were evaluated by measuring two perpendicular diameters of the tumor with a caliper. Individual tumor volume was calculated as *(V) = a x b^2^/2*, with *a* being the major and *b* the minor diameter. Mice were sacrificed by cervical dislocation after 3 months from the injection. For each mouse, primary tumor, lungs, liver, brain, bone marrow, and renal and lumbar aortic lymph nodes were collected. Primary tumors were halved, and half of them were used to extract total RNA, while the other was fixed in 10% neutral-buffered formalin and embedded in paraffin blocks.

### RNAseq analysis and bioinformatic analysis

Total RNA from the explanted xenografts was isolated using TRIzol (Thermo Fisher Scientific, Inc.), and RNA integrity was analyzed with the RNA 6,000 Nano Chip on an Agilent 2100 Bioanalyzer (Agilent Technologies). rRNA was depleted using the RiboCop rRNA Depletion Kits (Lexogen), and RNA-Seq libraries were generated using the CORALL Total RNA-Seq Library Prep Kit (Lexogen) according to manufacturer’s instructions. Size and quality of the libraries were assessed using a BioAnalyzer High Sensitivity Chip (Agilent). All samples were normalized to 2 nM and pooled equimolar. The library pool was sequenced on the NextSeq500 (Illumina) with 1x75bp, with 21 to 31 mio reads per sample. Gene expression was calculated by means of gene annotations from ENSEMBL (version 98) for human GRCh38 genome assembly after aligning raw FASTQ files using STAR (2.7.0f) (STAR https://pubmed.ncbi.nlm.nih.gov/23104886/) with default parameters with option “--quantMode GeneCounts”. Differential gene expression analysis was performed using DESeq2 package (Love *et al*, 2014). Null variance of Wald test statistic output by DESeq2 was re-estimated using the R-package FDR-tool (Strimmer, 2008) to re-calculate *P*- values—and adjusted using Benjamini-Hochburg method—for the final list of differentially expressed genes.

Gene ontology enrichment analysis was performed on the obtained gene list using topGO (Alexa *&* Rahnenfuhrer, 2019). Ten non-DE-genes having similar mean expression level per each DE-gene were selected for enrichment analysis as background using R package genefilter (https://www.rdocumentation.org/packages/genefilter/versions/1.54.2). GO plots for enrichment of biological process were plotted using R package ClusterProfiler (Yu *et al*, 2012). The list of DEGs involved in the top 10 GO terms was further analyzed by GSEA version 3.0 software (Broad Institute). Preranked analyses were performed by sorting differentially expressed genes according to their fold changes without prior filtering on significance or effect size. Enrichment map was used for visualization of the GSEA results. One thousand permutations were performed for each analysis. Disease and function bioinformatics analysis were generated through the use of IPA (QIAGEN Inc., https://www.qiagenbioinformatics.com/products/ingenuitypathway-analysis). The score is generated based on hypergeometric distribution, where the negative logarithm of the significance level is obtained by Fisher’s exact test at the right tail. The −log (*P*-value) > 2 was adopted as threshold, the Z-score > 2 was defined as the threshold of significant activation, while the Z-score < −2 was defined as the threshold of significant inhibition.

### Tube formation assay

The *in vitro* tube formation assay was performed as previously described (Hetheridge *et al*, 2011). Briefly, 3 x 10^4^ CAFs or HFFs were seeded on 10 mm glass coverslips (Thermo Fisher Scientific, Inc.) and, when they reached confluence, covered by a layer of HDBECs (1 x 10^4^). The medium (ECGM-2 w/o VEGF) was replenished every other day. When the cells were treated with APA or AT_1_R pharmacological inhibitors, the medium was refreshed 24 h after HDBEC plating and supplemented with the indicated concentration of amastatin (A1276; Sigma-Aldrich), candesartan (CV11974; Tocris), or vehicle (DMSO or MeOH respectively) every other day. After 10 days of co-culture, cells were fixed with 4% PFA/PBS for 10 min at 4°C, washed with PBS, and permeabilized with 0.05% Tween 20 in PBS. Samples were blocked with 3% BSA in PBS before incubation with the primary antibody αCD31 (BD Pharmingen). Images were acquired using SP5 confocal microscope (Leica). The mean tubule length was quantified using Fiji software (Schindelin *et al*., 2012). Tubule density and total tubule length were quantified using WimTube software program (Wimasis, Munich, Germany).

### Cell viability assay

Cells were seeded at a density of 1 x 10^4^/well in a 96-well culture plate for 24 h and then treated with the indicated concentrations of amastatin, candesartan, or vehicle (MeOH or DMSO, respectively). Forty-eight h later, cell viability was determined using the 3-(4,5-dimethylthiazol-2- yl)-2,5-diphenyltetrazolium bromide (MTT) (Sigma-Aldrich, Milan, Italy) according to the manufacturer’s instruction.

### Statistical analysis

All statistical tests were performed using GraphPad Prism version 7.00 for Mac (GraphPad Software). The data are presented as means ± standard deviations (SD). For comparisons consisting of two groups, means were compared by two-tailed Student’s t test. For comparison among three or more groups, means were compared using one-way or two-way ANOVA with Bonferroni posttest. Differences were considered statistically significant at a *P*-value of < 0.05.

## Acknowledgments

We thank Marcello Arsura for critically reviewing the manuscript and Bernd Zobiak and Antonio Virgilio Failla, HEXT, UKE Microscopic Imaging Facility (UMIF) for their help with the confocal microscopy. We thank Kerstin Reumann and Christina Herrde for excellent technical support with the NGS library preparation and sequencing. We are very grateful to “Rete Oncologica Piemontese” for providing the MCC specimens. The authors gratefully acknowledge Michela Salvo from the Histology Research Core Facility at University of Piemonte Orientale (Novara, Italy) for technical support in tissue processing and histological analysis and Valeria Caneparo at CAAD - Center for Translational Research on Autoimmune and Allergic Disease (Novara, Italy) for assistance with ddPCR assays.

This work was supported by grants from the Italian Ministry for University and Research—MIUR (PRIN 2017) to C.B., the AGING Project—Department of Excellence—DIMET to M.G., and the Università del Piemonte Orientale, and “Rete Oncologica Piemontese” to R.B. and M.G.. S.A. was supported by the Rotary Global Grant (2019-2020), which was sponsored by the Rotary Club of Novara and financed by the Rotary Foundation and University of Piemonte Orientale, Department of Excellence - DIMET, and by the Erich und Gertrud Roggenbuck Foundation, Hamburg (2020- 2021).

## Author contribution

Conception and design: SA; NF; MG. Development and methodology: SA; LM; CB; NF; MG. Acquisition of data: SA; LM; CB; SV; DI; GG; FC; NF; MG. Analysis and interpretation of data: SA; LM; CB; SV; DI; ILC; RB; NF; MG. Administrative, technical, or material support: SA; SV; DI; ILC; GG; FC; RB; NF; MG. Writing, review, and/or revision of the manuscript: SA; LM; CB; NF; MG. Final approval: All authors.

## Conflict of Interest

The authors declare no conflict of interest.

## The paper explained

### Problem

Merkel cell carcinoma (MCC) is a highly aggressive skin cancer, with most patients presenting with already advanced tumor stages at the time of diagnosis. The 5-year survival rate significantly drops with the presence of metastases. While immune therapy shows partial regression of MCC in the majority of cases, a significant proportion of patients develop immunotherapy-resistant tumor cells and even hyperprogressive disease courses. Analyzing the tumor microenvironment (TME), in particular the contribution of cancer-associated fibroblasts (CAFs)—the major component of TME— will significantly contribute to our understanding of MCC progression and might be helpful to identify new therapeutic targets.

### Results

This study describes the molecular and functional characterization of CAFs in MCC. CAFs were isolated from primary or recurrent MCC tumors and functionally characterized by gene expression and contractility assays. Their impact on MCC tumorigenesis was investigated in a xenograft mouse model. These experiments, together with transcriptome analyses of the xenografts obtained by subcutaneous co-injection of patient-derived CAFs and human MCC MKL-1 cells, demonstrate that the growth-promoting activity of CAFs is mediated by the APA/Ang II-III/AT_1_R axis, with expression of aminopeptidase A (APA) being the upstream triggering event.

### Impact

In this study, we have characterized, for the first time, the role of CAFS in MCC tumor progression. We demonstrate that those CAFs that exhibit tumor-promoting activity display high expression levels of aminopeptidase A (APA). Furthermore, we show a positive correlation of APA expression with CAF localization around blood vessels and enhanced tumor angiogenesis. These findings decipher APA and the APA/Ang II-III/ATR_1_R axis as a putative signaling pathway amenable to therapeutic intervention, especially since there are known inhibitors of this pathway already approved for other diseases.

## Expanded View Figure Legends

**Figure EV1. Characterization of the stromal components.** Scatter plots showing the number of cells per mm^2^ that were αSMA-1^+^ (green), CD8^+^ (red), and CD68^+^ (blue) in the tumor edges (A) and nests (B). The number of cells/mm^2^ was quantified from 10 independent tumor areas (represented as individual points) using the fast cell count tool by QuPath.

**Figure EV2. Tumor growth assessment**. Tumor growth curve of each mice group co-injected with MKL-1 and CAFs (rPt2 to pPt9) compared to mice injected with MKL-1 cells alone (black lines) or with MKL-1 plus HFFs (grey lines). Values at each time point are represented as mean ± SD within the mouse replicates for each group. Differences between curves were evaluated by two-way ANOVA. Statistical analysis at each time point was performed by two-way ANOVA followed post-hoc Bonferroni correction. \**P* < 0.05, \*\**P* < 0.01, \*\*\**P* < 0.001 *vs* MKL-1/HFF tumors or ^#^*P* < 0.05, ^##^*P* < 0.01, ^###^*P* < 0.001 *vs* MKL-1 tumors.

**Figure EV3. Ingenuity pathway analysis.** This analysis shows different diseases and functions predicted to be activated (z-score > 2, orange bars) or inhibited (z-score < 2, blue bars) in MKL- 1/CAFs *vs* MKL-1/CTRLs.

**Figure EV4. CAFs support angiogenesis in an *in vitro* model of tubulogenesis.** CAFs or HFF were co-cultured with HDBEC for 10 days. Representative images of CD31 immunostaining of the tubule-like structures (red). Nuclei were counterstained with DAPI (blue). Scale bars 100 μm.

**Figure EV5. Lack of enhanced or synergistic effects upon co-treatment with amastatin and candesartan.** rPt5 CAFs were co-cultured with HDBECs for 10 days in the presence of amastatin, candesartan, the combination of both inhibitors, or vehicle alone (MeOH for amastatin and DMSO for candesartan, respectively) at the indicated concentrations. Panel A shows representative images of CD31 immunostaining of the tubule-like structures (red). Nuclei were counterstained with DAPI (blue). Scale bars 100 μm. Panel B shows the quantification of mean tubule length using Fiji software. Panel C shows the quantification of the total tubule length as measured by WimTube software. Panel D shows the tubule density as measured by WimTube software. Statistical analysis was performed by classical one-way ANOVA followed by Bonferroni correction. \**P* < 0.05, \*\**P* < 0.01, \*\*\**P* < 0.001 *vs* vehicle control.

